# From Genes to Brains: Molecular Evolution and Mammalian Brain Cellular Diversity

**DOI:** 10.64898/2026.09.16.752041

**Authors:** Alexander Solovyov, Maksim Kazanskii, Alibek Suraganov, Artem Kasianov

## Abstract

Mammalian brains exhibit extensive diversity in size, cellular composition, and organization, yet the molecular evolutionary changes associated with this phenotypic diversification remain incompletely understood. Here, we investigated whether protein-sequence evolution covaries with quantitative differences in brain phenotypes across mammals. We integrated 29 brain and body traits with orthologous protein sequences from 18 mammalian species and obtained gene-specific molecular evolutionary measures for 10,395 genes. To reduce redundancy among strongly correlated phenotypes, we selected five representative traits that explained 88.12% of the standardized trait variance under linear reconstruction. Using RERconverge, we tested for associations between gene-specific relative evolutionary rates and evolutionary changes in these five representative traits. The primary analysis, based on log-transformed phenotypes, identified 25 significant gene–trait associations involving 23 genes, 19 of which were also recovered using untransformed phenotypes, indicating that a substantial subset of the associations was robust to phenotype transformation. Expression profiling of these shared candidate genes across adult human GTEx brain tissues revealed heterogeneous patterns ranging from broad expression across the examined tissues to low adult brain expression. Functional enrichment analysis further showed that the gene set retained for comparative analysis was enriched for diverse biological processes, including metabolic, cellular, and neural pathways. These results identify gene-specific evolutionary-rate associations with quantitative mammalian phenotypes and provide functional context for the genes represented in the comparative analysis. Our study provides a comparative framework linking relative evolutionary rates of protein-coding genes to quantitative variation in mammalian brain cellular architecture while accounting for shared evolutionary history.

## Introduction

Mammalian brains exhibit remarkable diversity in size, cellular composition, and internal organization. Across species, brain structures vary not only in mass but also in the numbers and densities of neurons and non-neuronal cells and in their distribution among major regions such as the cerebral cortex and cerebellum. These characteristics do not scale uniformly across mammals. For example, primate brain structures scale approximately isometrically with neuronal number, whereas rodent cerebral cortices increase in mass more rapidly than in neuronal number; insectivores combine rodent-like cortical scaling with primate-like cerebellar scaling [Herculano-Houzel et al., 2007, Sarko et al., 2009, Herculano-Houzel, 2012]. Consequently, similarly sized brains from different mammalian groups can contain markedly different numbers and densities of neurons, demonstrating that brain mass alone provides an incomplete description of brain cellular organization [Azevedo et al., 2009, Herculano-Houzel, 2012].

Mammalian brain diversification therefore encompasses multiple dimensions of phenotypic change, including neuronal abundance, density, cellular composition, and the allocation of cells among brain structures. These dimensions can evolve differently even among species with large brains. For example, the elephant cerebellum contains an exceptionally large fraction of the brain’s neurons, whereas its cerebral cortex contains fewer neurons than the human cerebral cortex despite its substantially greater mass [Herculano-Houzel et al., 2014]. Mammalian brain evolution thus reflects changes in cellular organization beyond a single axis of brain size, raising a central question: what molecular evolutionary changes accompany the diversification of brain cellular architecture across mammals?

The development and organization of the mammalian brain depend on gene networks involved in neurogenesis, neuronal differentiation, axon guidance, synaptic function, myelination, cell adhesion, and developmental regulation. Comparative studies have implicated evolutionary changes in protein-coding sequences, gene duplication, regulatory elements, and gene-expression programs in the diversification of neural phenotypes [Khaitovich et al., 2006, Gallego Romero et al., 2012, Rogers and Gibbs, 2014]. In primates, lineage-specific changes in brain gene expression, protein evolution, and regulatory sequences have been particularly well documented [Khaitovich et al., 2006, Somel et al., 2013, Shao et al., 2023, Zhuang et al., 2023]. These studies demonstrate that brain evolution involves changes across multiple levels of genome organization and regulation. However, much of this literature has focused on particular genes, mechanisms, or lineages—especially those associated with human and primate brain evolution—rather than asking on a genome-wide scale whether molecular divergence among species systematically tracks quantitative differences in brain cellular phenotypes. Relative evolutionary rate analysis provides a framework for addressing this question by testing whether lineage-specific changes in the evolutionary rates of protein-coding genes covary with evolutionary changes in quantitative phenotypes. This approach enables gene– phenotype relationships to be evaluated systematically across large numbers of genes and traits while incorporating the phylogenetic structure of the sampled species.

A central challenge in cross-species analyses is that species cannot be treated as statistically independent observations because their molecular and phenotypic characteristics are structured by shared evolutionary history [Felsenstein, 1985, Harvey and Pagel, 1991]. Closely related species generally share greater genetic similarity and may also resemble one another in morphological and physiological traits simply as a consequence of common ancestry. Consequently, an observed association between molecular divergence and differences in brain morphology may arise partly from their mutual dependence on phylogenetic relatedness rather than from a specific relationship between the evolution of a gene and the evolution of a brain phenotype. Phylogenetic comparative methods address this problem by explicitly incorporating information about evolutionary relationships among species when evaluating associations between biological variables [Felsenstein, 1985, Garland and Ives, 2000].

Expression data provide an additional means of characterizing the potential biological relevance of genes identified through comparative evolutionary analyses. Because genes associated with brain evolution may differ in their expression across neural tissues, examining the brain-expression profiles of candidate genes can help contextualize evolutionary-rate associations and identify whether particular candidates are broadly or regionally expressed in the adult human brain.

Here, we investigate whether molecular evolution in protein-coding genes across mammals is associated with quantitative divergence in brain cellular organization. We integrate comparative neuroanatomical measurements with orthologous protein sequences across 18 mammalian species, analyzing 10,395 genes and 29 morphological and cellular traits, including brain phenotypes and body mass. Because many of these traits are strongly correlated, we select five representative phenotypes for the primary gene-level analyses. Using RERconverge, we test whether gene-specific relative evolutionary rates across the mammalian phylogeny are associated with evolutionary changes in these representative phenotypes. We further characterize the brain-expression profiles of genes identified by RERconverge and examine the functional composition of the gene set retained for comparative analysis. By combining lineage-specific relative evolutionary rates, quantitative brain phenotypes, and phylogenetic information within a common comparative framework, this study provides a genome-wide assessment of the molecular evolutionary correlates of mammalian brain cellular diversification.

## Methods

### Brain and body morphological data and species set

Brain morphological data were obtained from the dataset of Herculano-Houzel et al. [2015]. The source dataset includes species-level measurements of brain structure mass, neuronal and non-neuronal cell numbers, cell densities, and related traits across mammalian species. We used all traits reported in Tables 1, 2, 3, 5, and 6, excluding traits related to the olfactory bulb. In total, 29 brain and body-related morphological traits were retained.

The initial species set was defined by the species present in the source dataset. For downstream analyses, we retained species for which orthologous protein sequences of human genes could be retrieved from OrthoDB [Tegenfeldt et al., 2025]. The final comparative dataset included 18 mammalian species: Homo sapiens, Mus musculus, Heterocephalus glaber, Condylura cristata, Mesocricetus auratus, Rattus norvegicus, Microcebus murinus, Cavia porcellus, Sciurus carolinensis, Oryctolagus cuniculus, Callithrix jacchus, Otolemur garnettii, Procavia capensis, Macaca fascicularis, Hydrochoerus hydrochaeris, Macaca mulatta, Tragelaphus strepsiceros, and Loxodonta africana. Species names matched exactly between the Herculano-Houzel et al. dataset and OrthoDB.

Hierarchical clustering of the 29 retained morphological traits revealed a pronounced block structure in the trait correlation matrix (Fig. 1). Most size-related variables, including brain-structure masses and neuronal and nonneuronal cell counts for the whole brain, cerebral cortex, cerebellum, and rest of brain, formed a large positively correlated cluster. Within this cluster, many pairwise Pearson correlations were close to one, indicating strong covariation among absolute brain size and cell-number traits across species.

**Figure 1.**
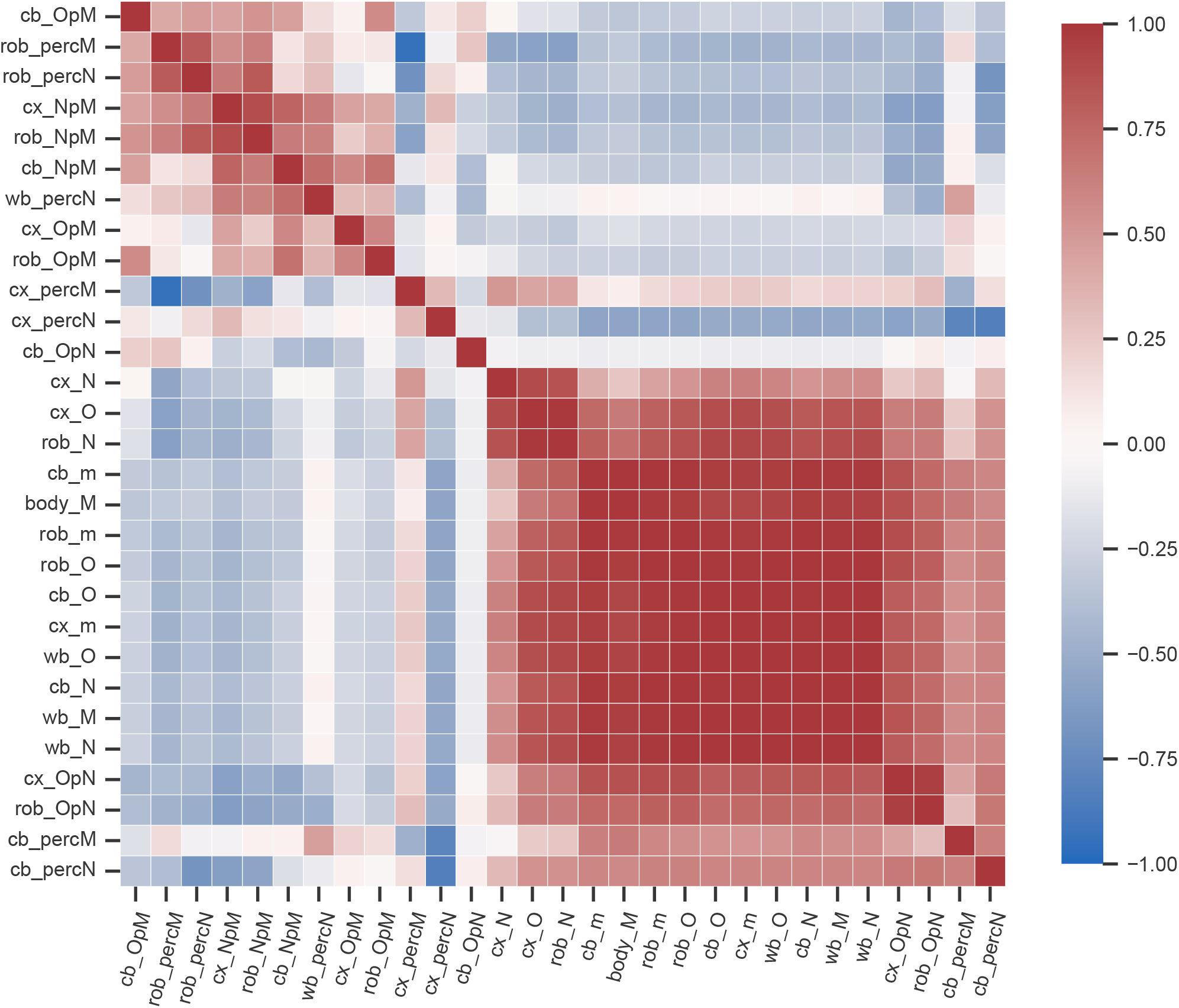
Hierarchically clustered Pearson correlation heatmap of the 29 retained brain and body-related morphological traits. Cell colors indicate Pearson correlation coefficients calculated across the final set of mammalian species.

In contrast, ratio- and percentage-based traits formed more separate clusters and often showed weaker or negative correlations with the absolute size-related traits. In particular, traits describing cell numbers per unit mass and neuronal or nonneuronal proportions were separated from the main size-related block. This clustering pattern indicates that the retained morphological traits capture both a dominant axis of overall brain and body size variation and additional relative-composition axes that are less directly aligned with absolute brain size.

### Selection of representative morphological traits

Many of the retained morphological traits were strongly correlated, therefore we reduced the trait set before the final gene–trait association analysis. All 29 traits were standardized to zero mean and unit variance. We then performed an exhaustive subset-selection analysis to evaluate how well smaller subsets of the original traits could represent the full 29-trait morphological space. For each subset size *k* = 1, …, 8, all possible combinations of *k* traits were evaluated.

For a given subset *S*, all 29 standardized traits were linearly reconstructed from the traits included in *S*. The total explained variation was calculated as

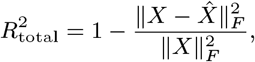

where *X* is the standardized matrix of the 29 morphological traits and 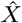 is its linear reconstruction from the selected subset. Because all traits were standardized, this value is equivalent to the mean *R*^2^ across the 29 traits. We also calculated the explained variation for the non-selected traits only, to assess how well each subset represented traits that were not themselves included in the predictor set.

The number of representative traits was chosen based on the relationship between subset size and explained variation. Explained variation increased rapidly for small values of *k*, whereas after *k* = 5, adding further traits produced progressively smaller gains relative to the increase in the number of downstream gene–trait tests. We therefore selected five traits as a compact representation of the original morphological trait set.

After choosing *k* = 5, we examined five-trait subsets with high total explained variation and prioritized subsets with lower internal redundancy. Internal redundancy was quantified using the mean and maximum absolute pairwise Pearson correlations among the selected traits. The final representative set consisted of *cx_N, cb_N pM, body_M, cx_percM*, and *cb_percN*. This subset explained 88.12% of the total variation across the 29 traits and 85.64% of the variation among the remaining non-selected traits. The mean absolute pairwise correlation within the selected subset was 0.255, and the maximum absolute pairwise correlation was 0.576. The pairwise correlation structure among the five selected representative traits is shown in Fig. 2.

**Figure 2.**
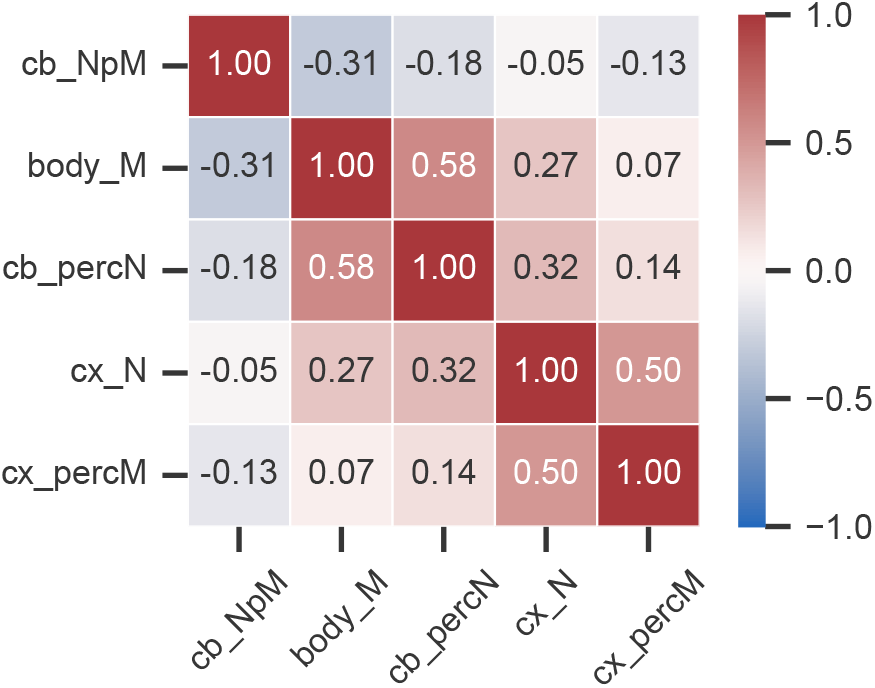
Pairwise Pearson correlations among the five representative morphological traits selected for the final analysis.

The final relative evolutionary rate analysis was restricted to these five representative morphological traits.

The traits were strictly positive and showed highly uneven, right-skewed distributions across species, especially for mass- and cell-number-related variables (Fig. 3); therefore, log transformation was used to reduce the influence of extreme values and to analyze proportional rather than absolute evolutionary changes.

**Figure 3.**
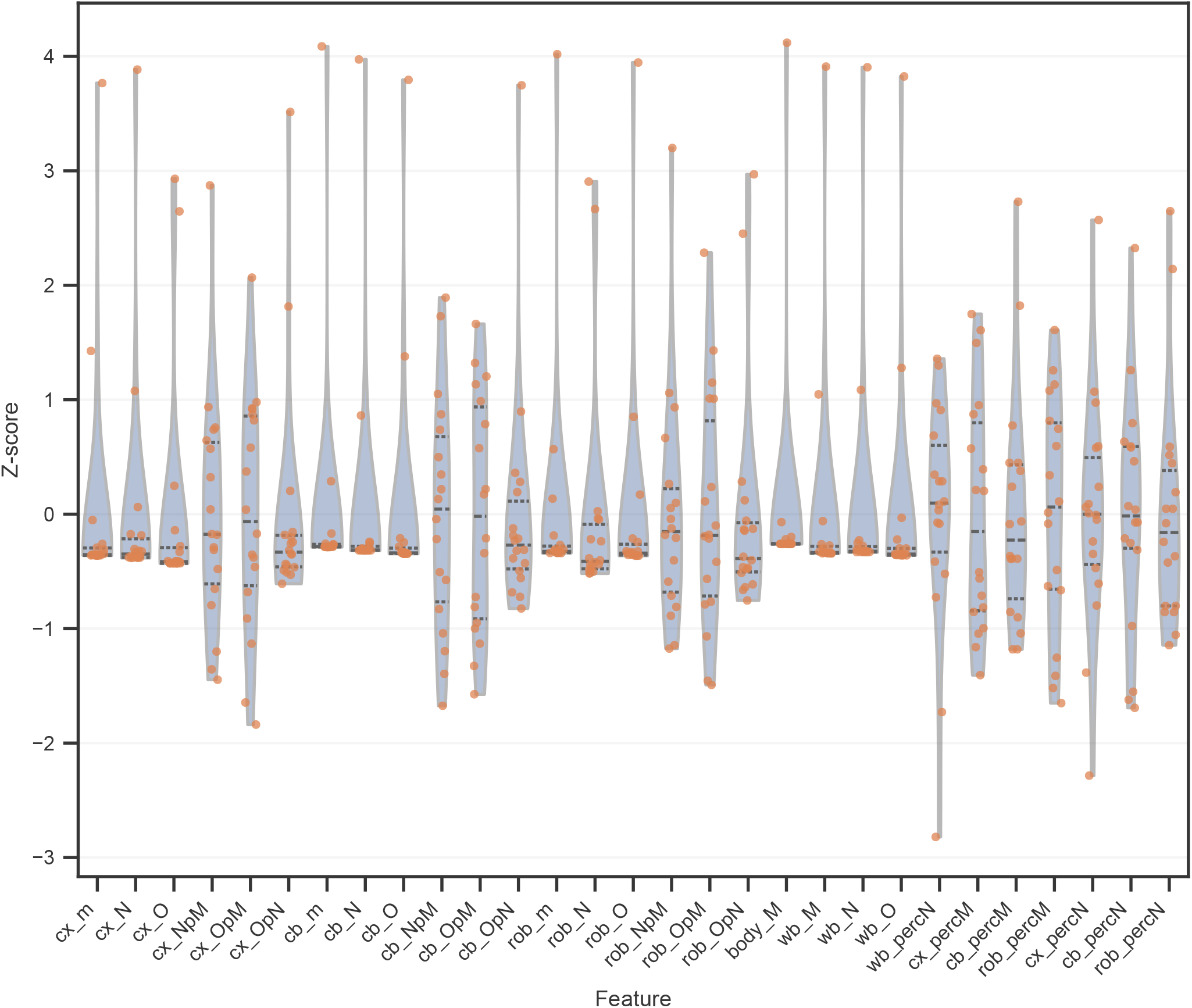
Distributions of standardized values for the 29 retained brain and body-related morphological traits.

### Ortholog identification and protein sequence retrieval

Orthologous protein sequences were retrieved from OrthoDB v12.2. The species set was first matched to OrthoDB using exact matches between the scientific names reported in the source brain morphology dataset and species names in the OrthoDB species table. Among the 40 species considered from the source dataset, 18 species had exact matches in OrthoDB, including *Homo sapiens*. Because human genes were used as the reference gene set, *Homo sapiens* was excluded from the non-human target species list for ortholog retrieval. The final target set therefore consisted of 17 non-human mammalian species with exact OrthoDB species-name matches.

Ortholog searches were performed at the Mammalian level using NCBI TaxID 40674. Human genes were mapped to OrthoDB human gene identifiers using Ensembl gene IDs. Version suffixes were removed from the input Ensembl gene IDs before matching them to Ensembl identifiers in the OrthoDB gene table. Human OrthoDB genes were then assigned to Mammalian-level orthologous groups, and all genes from these orthologous groups were retrieved for the 17 selected non-human mammalian species. Protein FASTA sequences for the resulting OrthoDB gene IDs were extracted from the OrthoDB amino acid FASTA file. Human protein sequences were also obtained from OrthoDB and included together with the non-human orthologous sequences.

For each human gene and species, a single representative protein sequence was retained. If multiple candidate sequences were available for a species, candidates were ranked by sequence quality and length: sequences with available FASTA records were prioritized, followed by sequences with fewer internal stop codons, fewer ambiguous or rare amino acid symbols, and greater sequence length. The OrthoDB gene identifier was used as a deterministic tie-breaker. This procedure produced one representative sequence per species for each retained human gene.

After representative-sequence selection, 11,829 human genes had an orthologous protein sequence available in all 17 selected non-human mammalian species. These genes were retained for downstream analysis. Together with the corresponding human sequence, each retained gene therefore had protein sequences for all 18 species in the comparative dataset. The 11,829 complete-coverage gene sets were carried forward to multiple sequence alignment and phylogenetic analysis.

To summarize ortholog availability in OrthoDB across species, we visualized the distribution of the number of species in which orthologs were identified for the queried human genes (Fig. 4).

**Figure 4.**
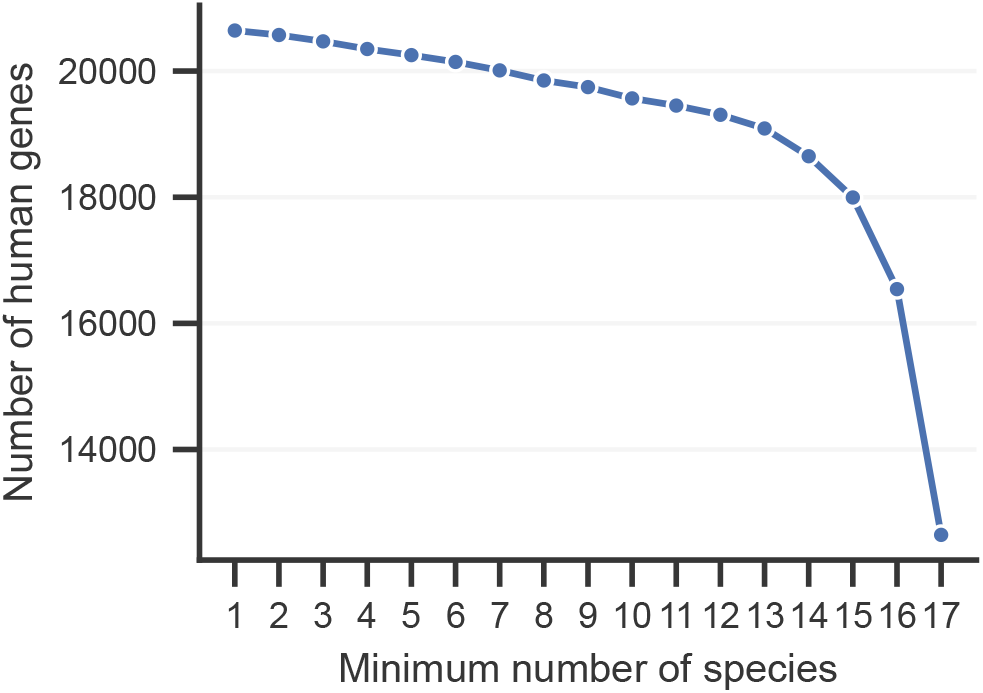
Cumulative distribution of human genes by the number of species with orthologs. The plot shows the number of human genes with orthologs in at least N species. The maximum observed ortholog coverage corresponded to 17 non-human mammalian species.

### Multiple sequence alignment

For each retained gene, amino acid sequences from Homo sapiens and the 17 non-human mammalian species were aligned using MUSCLE v5.3 [Edgar, 2022] with default parameters.

### Relative evolutionary rate analysis

We used RERconverge [Kowalczyk et al., 2019, Partha et al., 2019] to test whether gene-specific relative evolutionary rates were associated with evolutionary changes in the five representative brain morphological traits selected for the final analysis. A mammalian consensus tree obtained from the VertLife/MamPhy [Upham et al., 2019] phylogeny was pruned to the 18 species included in the comparative dataset and used to define a common species-tree topology. The temporal branch lengths of the VertLife tree were not used for estimation of gene-specific molecular evolutionary rates.

For each of the 10,395 protein alignments, gene-specific branch lengths were estimated with IQ-TREE v3.0.1 [Wong et al., 2026] while constraining the topology to the common 18-species tree (-te). Amino-acid substitution models were selected separately for each alignment using ModelFinder Plus (-m MFP) [Kalyaanamoorthy et al., 2017]. The resulting gene trees were required to contain the same 18 species, reproduce the same unrooted bipartitions, and have finite non-negative branch lengths.

To ensure consistent correspondence of branches among gene trees, gene-specific branch lengths were mapped by unrooted bipartitions onto a common reference representation. This procedure altered only the representation of the trees and was verified to preserve total tree length and all pairwise patristic distances. The resulting trees were imported into RERconverge using ‘readTrees()’.

Gene-specific RERs were then calculated using ‘getAllResiduals()’ with square-root transformation of molecular path lengths, weighted regression, a minimum species requirement of 18, and scale normalization. The resulting matrix contained 10,395 genes and 131 columns in the RERconverge phylogenetic-path representation. Because the number of finite RER values differed among genes, genes were considered testable only when at least 18 finite RER values were available. This criterion retained 10,031 genes for association testing.

For each of the five representative phenotypes *cb_N pM, body_M, cb_percN, cx_N*, and *cx_percM*, species-level trait values were converted to changes along the corresponding RERconverge phylogenetic paths using ‘char2Paths(metric = “diff”)’. The primary analysis was performed after natural-log transformation of the species-level phenotype values, whereas untransformed phenotypes were analyzed as a sensitivity analysis.

For each gene-trait pair, gene-specific RER values were correlated with phenotype-path values using Spearman rank correlation across finite paired paths. No winsorization was applied. Nominal P-values were obtained using R ‘cor.test(method = “spearman”, exact = FALSE)’. Multiple-testing correction was performed using the Benjamini-Hochberg procedure [Benjamini and Hochberg, 1995] across all finite tests involving the five selected traits. Associations with pooled five-trait FDR *q <* 0.05 were considered significant.

### Comparison of RER correlation profiles between highly brain-expressed genes and the remaining genes

To test whether genes with high expression in the human brain differed from the remaining genes in their relative evolutionary rate association profiles, we ranked genes by brain expression level using GTEx v11 data. For each gene, TPM values were summarized separately for each of the 13 GTEx brain tissues by calculating the median expression across samples within each tissue. We then calculated the mean of these 13 tissue-level median TPM values and used this value as a single brain expression score for each gene.

This analysis was restricted to genes that were testable in both the raw-scale and log-transformed RERconverge analyses. In total, 10,031 testable genes were included. Genes were sorted in descending order of the brain expression score, and the top 500 genes were defined as the highly brain-expressed group. The remaining 9,531 genes were used as the comparison group.

For each of the five representative morphological traits, we compared the distributions of Spearman correlation coefficients obtained from the RERconverge analyses between the highly brain-expressed genes and the remaining genes. Raw-scale and log-transformed RERconverge results were analyzed separately. All available Spearman correlation coefficients were included in this comparison, irrespective of the corresponding RERconverge nominal *p*-value or FDR value.

For each trait and each phenotype scale, the two distributions of Spearman correlation coefficients were compared using a two-sided two-sample Kolmogorov–Smirnov test implemented in SciPy [Virtanen et al., 2020]. The resulting *p*-values were adjusted across the five representative morphological traits using the Benjamini–Hochberg false discovery rate procedure. Traits with BH-adjusted *q <* 0.05 were considered to show significant distributional differences between the highly brain-expressed genes and the remaining genes.

To assess sensitivity to the choice of expression threshold, the comparison was repeated using the top 100, 200, 1,000, and 2,000 genes ranked by the same brain-expression score.

### KEGG pathway enrichment analysis

KEGG pathway enrichment analysis was performed to compare the 10,395 genes retained for comparative analysis with genes not included in the analyzed set. The comparison set was generated from the GTEx v11 gene-level expression table by extracting human gene symbols and removing all genes present in the analyzed set. After duplicate removal, this comparison set contained 62,926 unique gene symbols.

Enrichment was performed separately for the analyzed and comparison gene sets using the KEGG_2021_Human library and the local over-representation analysis implemented in GSEApy [Fang et al., 2023]. Input gene lists were restricted to genes represented in at least one KEGG pathway, and the enrichment background was defined as all unique genes present in the KEGG_2021_Human library. Pathways with adjusted *p <* 0.05 were considered significantly enriched. Significant pathways were then classified as enriched only in the analyzed gene set, enriched only in the comparison gene set, or shared between both gene sets.

## Results

### Relative evolutionary rate analysis identifies a stable core of genes associated with brain morphological traits

RERconverge analysis was performed to test whether gene-specific relative evolutionary rates were associated with evolutionary changes in the five representative morphological traits selected for the final analysis: *cb_N pM, body_M, cb_percN, cx_N*, and *cx_percM*. The primary analysis used log-transformed trait values, and an additional sensitivity analysis was performed using the original untransformed trait values.

In the log-transformed analysis, 25 gene–trait associations remained significant after Benjamini–Hochberg correction across all finite tests in the five-trait panel, involving 23 unique genes (Fig. 5). Significant associations were detected for all five representative traits, but their distribution was uneven. The largest number of significant associations was observed for the percentage of neurons in the cerebellum (*cb_percN*; 10 associations), followed by the percentage of cortical mass (*cx_percM*; 8 associations) and the number of cortical neurons (*cx_N*; 4 associations). Fewer associations were detected for cerebellar neurons per unit mass (*cb_N pM*; 2 associations) and body mass (*body_M*; 1 association).

**Figure 5.**
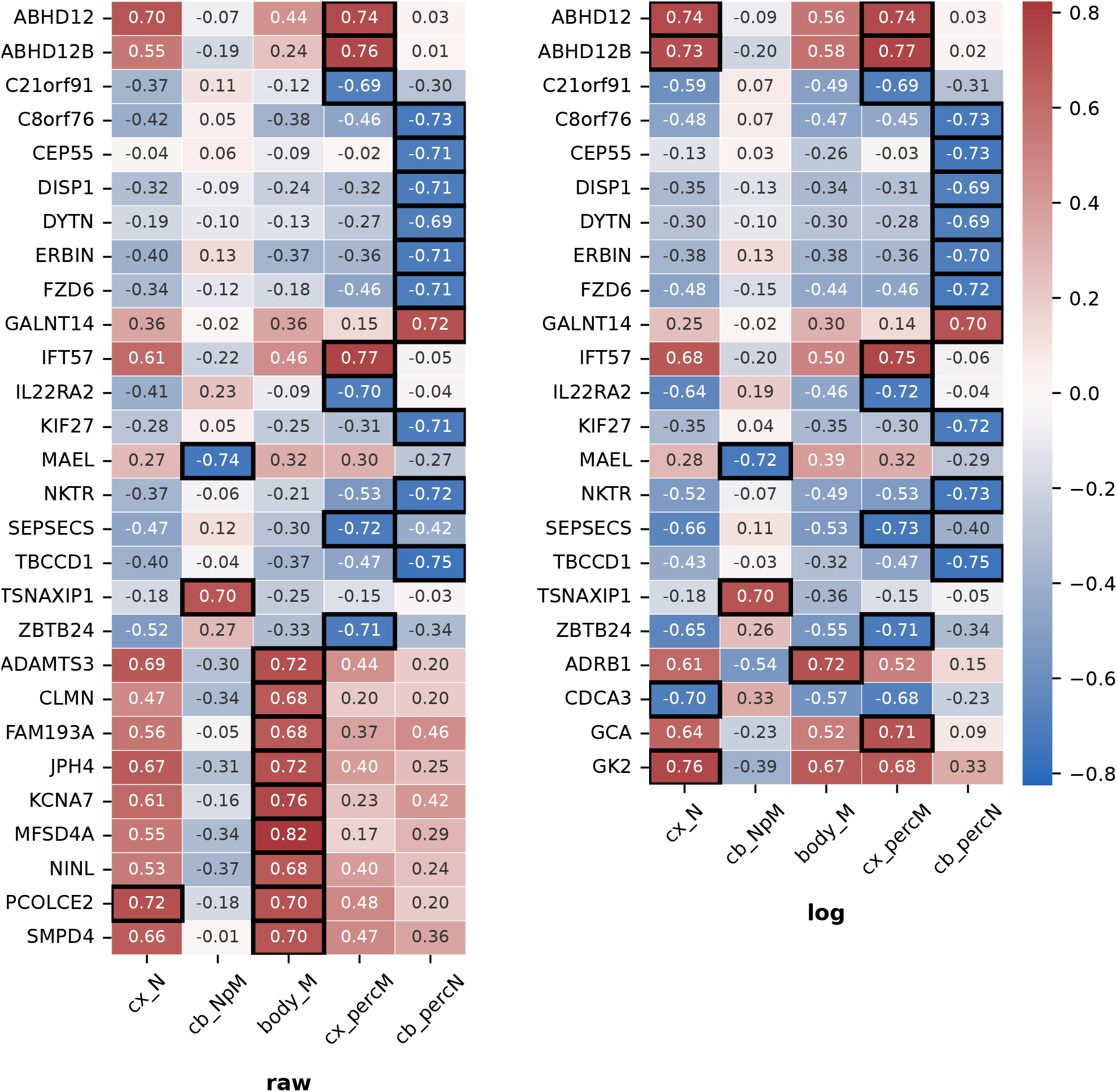
RERconverge associations between gene-specific relative evolutionary rates and the five representative morphological traits. The left heatmap shows results obtained using trait data on the original scale (raw), whereas the right heatmap shows results obtained after log transformation of trait values prior to the analysis (log). Rows show genes with at least one significant gene–trait association after Benjamini– Hochberg correction across all finite tests in the selected five-trait panel. Thick borders indicate associations with *q <* 0.05.

The sensitivity analysis based on untransformed trait values identified 29 significant gene–trait associations involving 28 unique genes. In this analysis, significant associations were again detected for all five traits, but the distribution across traits differed from the log-transformed analysis. The largest number of associations was observed for *cb_percN* and *body_M*, with 10 and 9 significant associations, respectively, followed by *cx_percM* with 7 associations. Fewer associations were detected for *cb_N pM* and *cx_N*, with 2 and 1 significant associations, respectively.

Most genes identified in the log-transformed analysis were also detected in the untransformed analysis. Nineteen genes were shared between the two analyses: *ABHD*12, *ABHD*12*B, C*21*orf* 91, *C*8*orf* 76, *CEP* 55, *DISP* 1, *DY TN, ERBIN, FZD*6, *GALNT* 14, *IFT* 57, *IL*22*RA*2, *KIF* 27, *M AEL, NKTR, SEPSECS, TBCCD*1, *TSN AXIP* 1, and *ZBTB*24. For these shared genes, the direction and magnitude of the correlations were broadly consistent between the raw-scale and log-transformed analyses, indicating that a substantial part of the RERconverge signal was robust to phenotype transformation.

The main difference between the two analyses concerned associations with *body_M*. Several genes were significant only in the untransformed analysis, including *ADAM TS*3, *CLMN, F AM* 193*A, JPH*4, *KCNA*7, *MFSD*4*A, NINL, PCOLCE*2, and *SMPD*4. These raw-scale-specific associations were mainly positive associations with *body_M*, suggesting that some body-mass-related RER associations are sensitive to the scale on which the phenotype is analyzed. Conversely, four genes were significant only in the log-transformed analysis: *ADRB*1, *CDCA*3, *GCA*, and *GK*2. This body-mass-specific sensitivity indicates that phenotype transformation can influence the detection of associations for traits with large absolute variation.

### Brain expression profiles of RERconverge candidate genes

To further characterize the genes identified by the RERconverge analysis, we examined GTEx expression profiles for genes that were significant in both the raw-scale and log-transformed analyses. This intersection contained 19 genes. Expression was summarized across 13 adult human brain tissues using log_2_(TPM + 1) values.

The candidate genes showed heterogeneous expression patterns across brain tissues (Fig. 6). Several genes were broadly expressed across most brain regions, including *ABHD*12, *NKTR, ERBIN, IFT* 57, *C*8*orf* 76, *TSN AXIP* 1, *TBCCD*1, and *FZD*6. Among these, *ABHD*12 and *NKTR* showed the highest overall expression, with consistently elevated expression across nearly all GTEx brain tissues. Other genes, including *C*21*orf* 91, *ZBTB*24, *SEPSECS, ABHD*12*B, DISP* 1, and *KIF* 27, showed lower but detectable expression across multiple brain regions.

**Figure 6.**
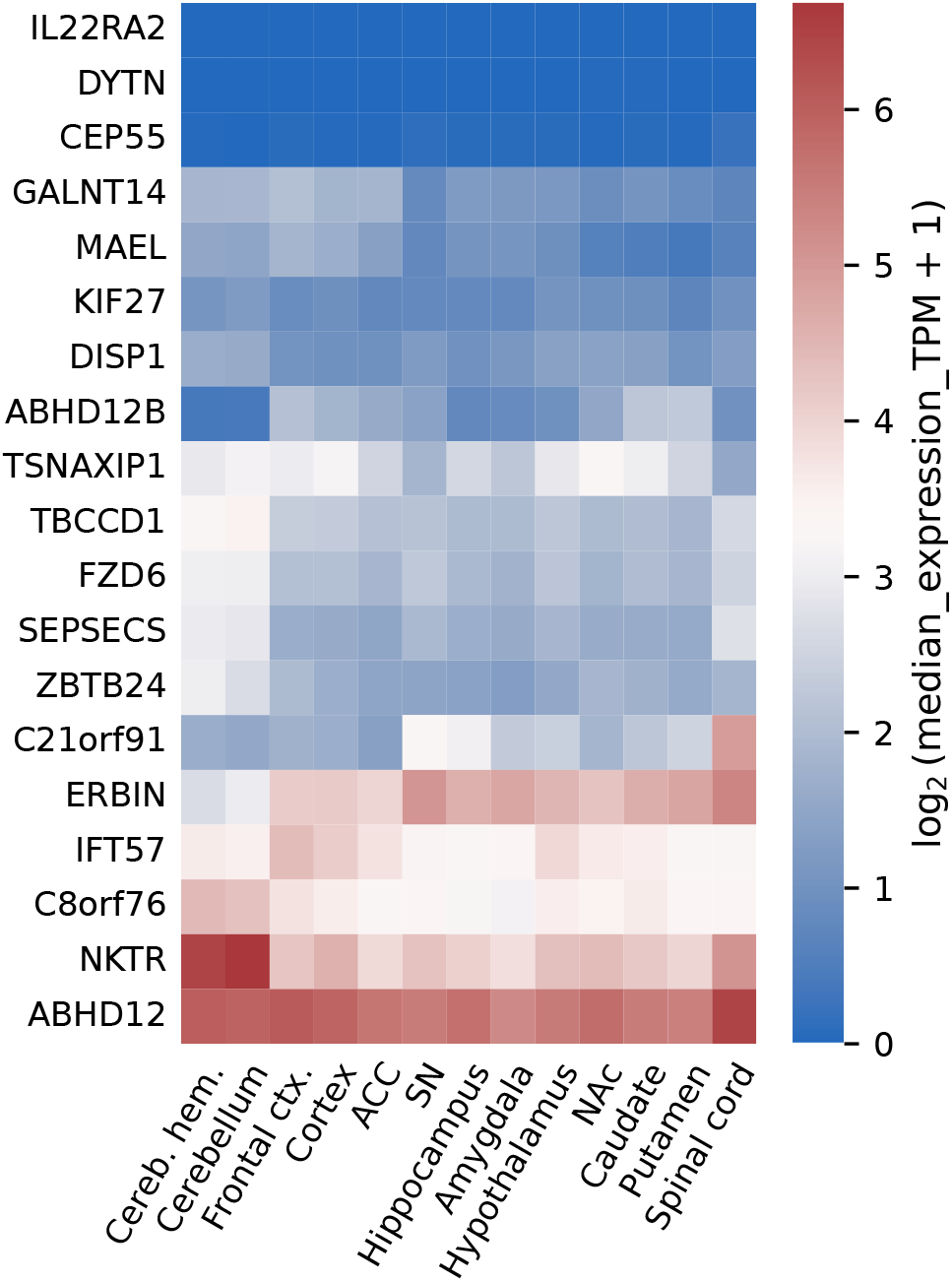
Brain expression profiles of genes significant in both raw-scale and log-transformed RERconverge analyses. Columns show GTEx brain tissues. Cell colors indicate log_2_(median TPM + 1), where median expression was calculated across samples within each tissue.

In contrast, a subset of RERconverge candidate genes showed low expression in adult brain tissues. *IL*22*RA*2, *DY TN*, and *CEP* 55 had expression values close to zero across most brain regions, indicating that not all genes with significant evolutionary-rate associations are highly expressed in the adult human brain. Thus, the RERconverge signal was not restricted to genes with uniformly high adult brain expression.

The sample-level expression distributions were consistent with the median-expression heatmap (Fig. 7). Broadly expressed genes such as *ABHD*12, *NKTR, ERBIN*, and *IFT* 57 showed detectable expression across many tissues and samples, whereas low-expression genes such as *IL*22*RA*2, *DY TN*, and *CEP* 55 had distributions concentrated near zero. Several genes also showed regional variation, including *C*21*orf* 91, *ABHD*12*B, GALN T* 14, *M AEL*, and *TSN AXIP* 1, suggesting that some candidate genes may have regionally heterogeneous expression profiles within the brain.

**Figure 7.**
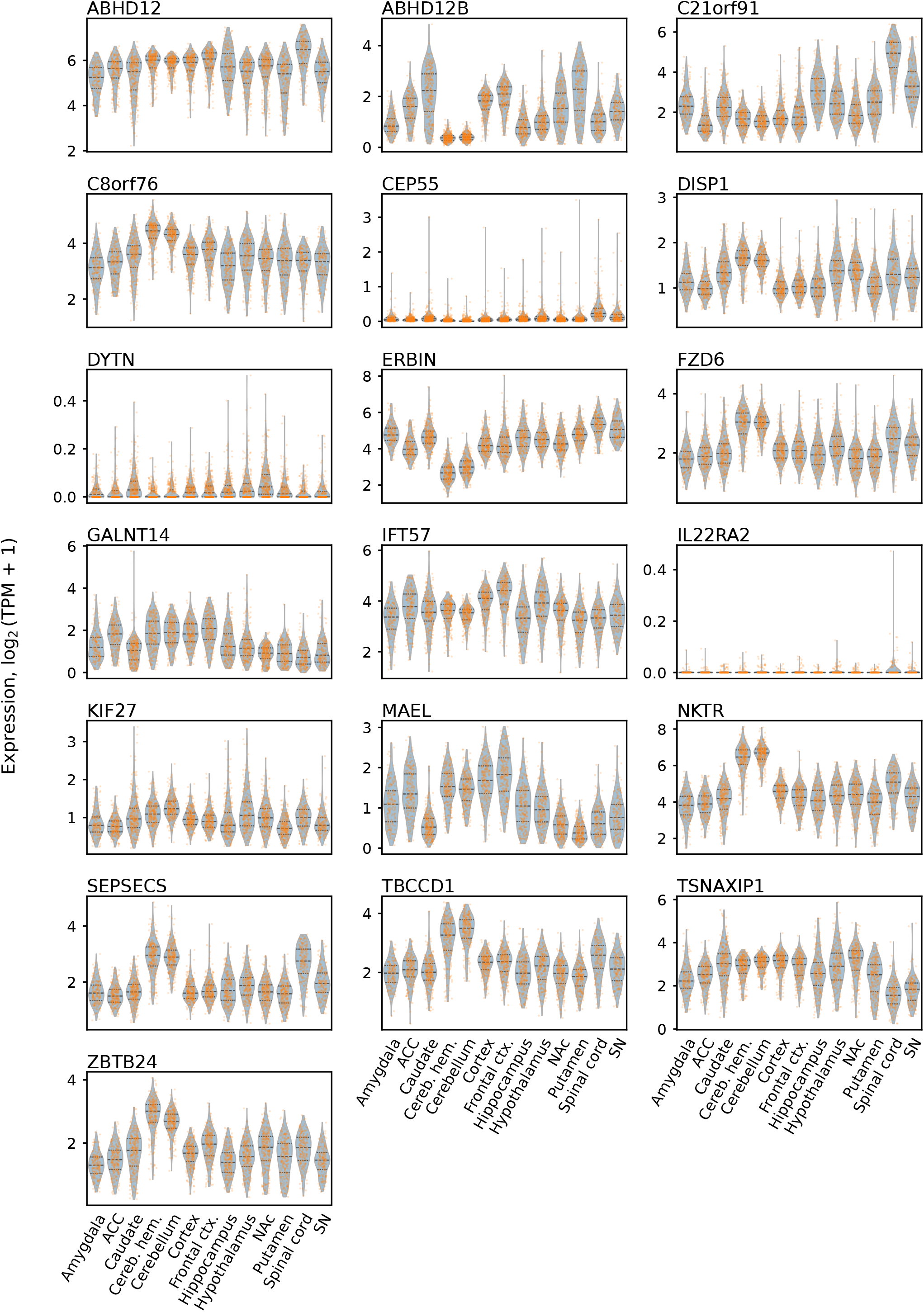
Sample-level expression distributions across GTEx brain tissues for genes significant in both raw-scale and log-transformed RERconverge analyses. Expression values are shown as log_2_(TPM + 1).

### Highly brain-expressed genes show distinct RER association profiles

We next examined whether genes with high expression in the human brain differed in their RERconverge correlation profiles. The analysis was restricted to 10,031 genes that were testable in both the raw-scale and log-transformed analyses. Genes were ranked by mean brain expression based on GTEx data, and the top 500 genes were compared with the remaining 9,531 genes. At this cutoff, three traits showed significant differences between the two distributions after Benjamini–Hochberg correction in both the raw-scale and log-transformed analyses: the number of cortical neurons (*cx_N*), the percentage of cortical mass (*cx_percM*), and the percentage of neurons in the cerebellum (*cb_percN*). For the raw-scale analysis, the corresponding KS statistics were *D* = 0.077 (*q* = 0.0105), *D* = 0.086 (*q* = 0.00429), and *D* = 0.102 (*q* = 4.75 *×* 10^*−*4^), respectively. For the log-transformed analysis, they were *D* = 0.095 (*q* = 8.72 *×* 10^*−*4^), *D* = 0.088 (*q* = 0.00213), and *D* = 0.112 (*q* = 6.67 *×* 10^*−*5^), respectively (Supplementary Fig. S1). For all three traits, the top 500 brain-expressed genes showed a modest shift toward more positive RERconverge correlation coefficients relative to the remaining genes.

We additionally repeated the analysis using the top 100, 200, 1,000, and 2,000 genes ranked by mean brain expression to assess sensitivity to the expression cutoff. No trait was significant in both phenotype transformations at the top-100 cutoff, whereas *cb_percN* was significant in both analyses at the top-200 cutoff. The same three traits identified with the top 500 genes (*cx_N, cx_percM*, and *cb_percN*) were also significant in both analyses at the top-1,000 cutoff. At the top-2,000 cutoff, all five examined traits were significant in both the raw-scale and log-transformed analyses.

### Functional enrichment of the analyzed gene set

To assess whether genes retained for comparative analysis differed from excluded genes in their functional representation, we compared KEGG pathway enrichment between the 10,395 analyzed genes and the genes excluded by filtering. Pathways with FDR-adjusted *P <* 0.05 were considered significantly enriched. Forty-six KEGG pathways were significantly enriched among the analyzed genes, compared with nine among the excluded genes, with no significantly enriched pathways shared between the two sets.

The analyzed gene set was enriched for multiple metabolic pathways, including the citrate cycle, fatty acid degradation, purine metabolism, glycolysis/gluconeogenesis, pyruvate metabolism, glutathione metabolism, amino acid metabolism, and steroid biosynthesis. Several pathways involved in glycan biosynthesis were also enriched, including N-glycan biosynthesis, mucin type O-glycan biosynthesis, and other types of O-glycan biosynthesis, together with pathways related to lysosomal function, ABC transporters, and cell adhesion molecules. In addition, axon guidance and neuroactive ligand–receptor interaction, which are associated with neural development and signaling, respectively, were significantly enriched among the analyzed genes.

By contrast, pathways significantly enriched among genes excluded from the comparative analysis included olfactory transduction, several immune- or disease-related pathways, ribosome, spliceosome, alcoholism, and circadian entrainment. This pattern may partly reflect the strict cross-species orthology filtering used in our analysis. Mammalian olfactory receptor repertoires undergo extensive lineage-specific gains, losses, and pseudogenization [Niimura and Nei, 2007], and such gene-family turnover can complicate orthology inference across divergent species [Tekaia, 2016]. These evolutionary dynamics could therefore contribute to the preferential exclusion of some olfactory and immune-related genes.

This explanation, however, is unlikely to apply uniformly to all pathways enriched among the excluded genes. Ribosomes possess a universally conserved evolutionary core across bacteria and eukaryotes [Melnikov et al., 2012], and key features of spliceosomal organization and dynamics are conserved between yeast and metazoans [Will and Lührmann, 2011]. Moreover, KEGG pathways defined by disease or infection labels represent molecular interaction networks and may include genes with diverse biological functions and evolutionary histories; they should therefore not be interpreted as consisting exclusively of rapidly evolving host-defense genes. Thus, the enrichment profile of the excluded gene set likely reflects multiple features of the filtering procedure and gene-set composition rather than providing direct evidence that all excluded pathways, or all genes assigned to them, are lineage-specific or rapidly evolving.

## Discussion

Understanding how molecular evolution relates to the diversification of complex phenotypes remains a central challenge in comparative biology. Here, we examined whether evolutionary changes in protein-coding genes covary with quantitative differences in mammalian brain cellular organization while accounting for shared evolutionary history. Starting from 29 morphological and cellular traits, including brain phenotypes and body mass, we selected five representative traits that captured most of the variation in the original morphological space while reducing redundancy among strongly correlated phenotypes. Using RERconverge, we then tested whether gene-specific deviations in evolutionary rate across the mammalian phylogeny were associated with evolutionary changes in these representative phenotypes.

The primary analysis using log-transformed phenotypes identified 25 significant gene–trait associations involving 23 genes, with significant associations detected for all five representative traits. Nineteen of these genes were also identified when the analysis was repeated using phenotypes on their original scale. The recovery of this substantial common set across the two phenotype representations indicates that many of the detected relative-rate associations are not dependent on a single transformation of the morphological data. At the same time, the differences between the two analyses demonstrate that phenotype representation can influence gene-level inference. This was particularly apparent for body mass, for which several additional associations were detected using untransformed values. Such sensitivity is plausible for phenotypes spanning large absolute ranges because differences on the original scale emphasize absolute evolutionary changes, whereas differences after logarithmic transformation more closely represent proportional changes. The overlap between the two analyses therefore provides evidence for associations that are comparatively robust to phenotype transformation while also emphasizing that the scale on which a quantitative phenotype is represented can affect the associations detected.

The distribution of significant associations across the five representative traits was heterogeneous. In the primary log-transformed analysis, the largest numbers of significant associations were detected for the percentage of neurons in the cerebellum and the percentage of cortical mass, whereas comparatively few associations were detected for body mass and cerebellar neurons per unit mass. This heterogeneity suggests that molecular evolutionary associations are not distributed uniformly across different dimensions of mammalian morphology. Brain cellular diversification encompasses multiple partially distinct features, including absolute neuronal numbers, cellular density, and the relative allocation of cells and mass among brain structures. The gene-level associations identified here are therefore more consistent with trait-dependent molecular correlates than with a single molecular axis underlying overall brain size or organization.

Molecular and phenotypic evolution can covary for several reasons. A gene may contribute directly to the development or function of the associated phenotype, but an association can also arise through correlated selection on interacting biological processes or through other lineage-specific evolutionary changes. The associations identified by RERconverge should therefore be interpreted as candidates for evolutionary coupling between molecular and phenotypic change rather than as evidence of causal genotype–phenotype relationships. Nevertheless, the recovery of a substantial common set of significant associations across the two phenotype transformations supports the broader conclusion that quantitative diversification of mammalian brain architecture is accompanied by detectable, trait-dependent patterns of protein evolution.

Expression profiling provided additional biological context for the genes recovered in both RERconverge analyses. The 19 shared candidate genes showed heterogeneous expression across the examined adult human GTEx brain tissues. Genes including *ABHD*12, *NKTR, ERBIN*, and *IFT* 57 were broadly expressed across the examined tissues, whereas other candidates, including *IL*22*RA*2, *DY TN*, and *CEP* 55, showed little expression in the adult human brain. Several candidates also exhibited variation among brain tissues. These observations indicate that evolutionary-rate associations with mammalian morphological phenotypes are not restricted to genes that are uniformly highly expressed in the adult human brain. However, the GTEx analysis is descriptive and should not be interpreted as demonstrating that adult human brain expression explains the evolutionary-rate associations. Expression during development, expression in particular cell types, or expression patterns in other mammalian species may provide different functional contexts for these genes.

The functional enrichment analysis also provides important context for the composition of the gene set available for comparative analysis. Requiring orthologous protein sequences across the complete 18-species set necessarily restricted the analysis to a subset of protein-coding genes. The 10,395 retained genes were enriched for a diverse collection of KEGG pathways, including pathways related to energy metabolism, cellular homeostasis, glycosylation, membrane and cellular organization, as well as axon guidance and neuroactive ligand– receptor interaction. By contrast, genes excluded from the comparative analysis showed enrichment for a different set of pathways, including olfactory, immune/infection-related, and general cellular categories. These results demonstrate that the complete-ortholog requirement produced a functionally non-random analyzed gene set and should therefore be considered when interpreting the scope of the gene-level results. This enrichment analysis characterizes the composition of the analyzed gene set rather than demonstrating that particular functional pathways have stronger evolutionary-rate associations with brain phenotypes.

The distributed and trait-dependent character of the gene-level results is consistent with a broader comparative literature showing that mammalian brain evolution involves coordinated changes across multiple levels of biological organization rather than variation in brain size alone. Recent analyses of brain– body scaling across large species samples have emphasized that macroevolutionary scaling relationships can emerge from heterogeneous evolutionary changes occurring within lineages [Baker et al., 2026]. At the cellular level, comparative analyses have likewise identified both conserved and lineage-dependent patterns. Hippocampal neuron numbers across mammals retain substantial signatures of phylogenetic history and ecology [Maliković et al., 2025], whereas glial densities and cell-type ratios show considerable conservation within homologous brain regions across mammalian lineages [Pinto-Duarte et al., 2025]. Together with earlier demonstrations of lineage-specific neuronal scaling rules [Herculano-Houzel et al., 2007, Sarko et al., 2009, Herculano-Houzel et al., 2015], these observations indicate that mammalian brain diversification reflects a combination of conserved cellular constraints and lineage-dependent changes in cellular composition.

Comparative genomic studies similarly suggest that substantial phenotypic diversification can coexist with conservation of many molecular components. Neuron-associated coding sequences and regulatory regions can remain strongly conserved across amniotes despite interspecific variation in neuronal cellular phenotypes [Xu and Herculano-Houzel, 2025]. At the same time, evolutionary changes in particular gene sets have been associated with the expansion of major brain structures. For example, comparative analyses of anthropoid primates have linked patterns of molecular selection in developmental genes to variation in neocortical and cerebellar expansion [Harrison and Montgomery, 2017]. More broadly, comparative studies have identified conserved scaling relationships in neuronal biophysics [Beaulieu-Laroche et al., 2021], synaptic density and connectivity [Sherwood et al., 2020], and large-scale cortical connectivity [van den Heuvel et al., 2016], despite pronounced differences in brain size and cellular composition among species. The gene-specific and trait-dependent associations observed here fit within this broader picture in which some components of mammalian neural systems remain constrained while others undergo lineage-dependent evolutionary change.

Accounting for phylogenetic relatedness is particularly important when interpreting cross-species molecular–phenotypic associations. Closely related species tend to resemble one another both molecularly and phenotypically because of shared ancestry, potentially generating apparent associations in the absence of phenotype-specific evolutionary coupling [Felsenstein, 1985, Harvey and Pagel, 1991]. RERconverge provides a phylogenetic framework by analyzing relative evolutionary rates and phenotype changes across paths on a common mammalian topology. This framework incorporates phylogenetic structure by evaluating molecular and phenotypic changes along corresponding evolutionary paths. Nevertheless, it cannot eliminate all possible effects of lineage-specific history, unmeasured species characteristics, or evolutionary processes unrelated to the phenotypes considered here. The detected associations should therefore be viewed as comparative evolutionary signals that generate hypotheses about gene– phenotype relationships rather than as direct evidence of functional or causal mechanisms.

Several additional limitations should be considered. First, the comparative dataset contained only 18 mammalian species. Although these species span multiple mammalian lineages, the sample remains small relative to mammalian diversity and limits statistical power as well as the ability to distinguish lineage-specific from broadly recurrent evolutionary patterns. Expanding the analysis to additional species with comparable cellular phenotypes would provide a stronger basis for evaluating the generality of the detected associations. Second, the analysis was restricted to protein-coding genes with orthologous sequences available across the complete species set. This requirement preferentially retains relatively well-conserved and broadly represented genes while excluding lineage-specific genes, gene gains and losses, and many rapidly evolving sequences. The functional enrichment differences between retained and excluded genes further demonstrate that this filtering step affects the biological composition of the analyzed gene set. Regulatory evolution, which has an important role in phenotypic diversification, is also not captured directly by protein-sequence analyses [Gallego Romero et al., 2012, Somel et al., 2013, Zhuang et al., 2023].

Third, the primary analysis used natural-log-transformed phenotypes, with the untransformed analysis serving as a sensitivity analysis. Although the substantial overlap between the two analyses is encouraging, the differences observed between them demonstrate that results can depend on phenotype representation. Alternative phenotype transformations and explicit models of continuous-trait evolution could therefore reveal additional relationships. Fourth, human GTEx data were used only to characterize the adult brain-expression profiles of RERconverge candidate genes. Adult human expression does not establish equivalent expression levels, tissue specificity, cell-type specificity, or developmental expression patterns across the other mammalian species included in the comparative analysis. Cross-species and developmental transcriptomic datasets would provide a more direct basis for investigating how expression evolution relates to the sequence-based associations identified here. Finally, despite multiple-testing correction, the gene-level associations arise from a comparatively small species set and should be validated in expanded comparative datasets and, where possible, using independent functional or evolutionary evidence.

Despite these limitations, the analysis identifies a set of protein-coding genes whose relative evolutionary rates covary with evolutionary changes in quantitative mammalian phenotypes. A substantial subset of these genes was recovered under both log-transformed and untransformed phenotype representations, while the differences between the two analyses demonstrate that some associations are sensitive to phenotype scale. The heterogeneous adult human brain-expression profiles of the shared candidates further indicate that these evolutionary associations are not restricted to genes with uniformly high adult brain expression. Together, these findings support a view in which the molecular correlates of mammalian brain diversification are gene-specific and trait-dependent, with their detection also influenced by the representation of quantitative phenotypes. Expanding this framework to larger species sets, regulatory sequence evolution, cross-species transcriptomic data, and additional phylogenetic models should help clarify how changes at different levels of genome organization contribute to the diversity of mammalian brain cellular architecture.

## Use of Large Language Models

A large language model (ChatGPT, OpenAI) was used during manuscript preparation for language polishing, including improvements to grammar, clarity, and wording. It was not used to generate or analyze data, perform statistical analyses, or draw scientific conclusions. All manuscript content was reviewed and approved by the authors.

## Data Availability

All primary data analyzed in this study were obtained from publicly available sources. Derived data generated during the analyses, including processed phenotype data, multiple sequence alignments, phylogenetic trees, and relative evolutionary rate matrices, are available in the accompanying GitHub repository.

## Code Availability

The code used for data processing, analysis, and figure generation is publicly available at: https://github.com/solovyov-alexander/Gene-brain-project.

## Author Contributions

Conceptualization: Maksim Kazanskii, Artem Kasianov;

Methodology: Alexander Solovyov;

Formal analysis: Alexander Solovyov;

Investigation: Alexander Solovyov, Alibek Suraganov;

Data curation: Alexander Solovyov, Maksim Kazanskii;

Visualization: Alexander Solovyov;

Validation: Alexander Solovyov;

Writing – original draft: Alexander Solovyov, Maksim Kazanskii;

Writing – review and editing: Alexander Solovyov, Maksim

Kazanskii, Alibek Suraganov, Artem Kasianov;

Supervision: Artem Kasianov;

Project administration: Maksim Kazanskii.

## References

Frederico A. C. Azevedo, Ludmila R. B. Carvalho, Lea T. Grinberg, José Marcelo Farfel, Renata E. L. Ferretti, Renata E. P. Leite, Wilson Jacob Filho, Roberto Lent, and Suzana Herculano-Houzel. Equal numbers of neuronal and nonneuronal cells make the human brain an isometrically scaled-up primate brain. J Comp Neurol, 513(5):532–541, 2009. doi: 10.1002/cne.21974.

Joanna Baker, Robert A. Barton, and Chris Venditti. Macroevolutionary brain scaling is a microevolutionary metaphenomenon. Nature Communications, 17:136, 2026. doi: 10.1038/s41467-025-66843-0.

Lou Beaulieu-Laroche, Norma J. Brown, Marissa Hansen, Enrique H. S. Toloza, Jitendra Sharma, Ziv M. Williams, Matthew P. Frosch, Garth Rees Cosgrove, Sydney S. Cash, and Mark T. Harnett. Allometric rules for mammalian cortical layer 5 neuron biophysics. Nature, 600(7888):274–278, 2021. doi: 10.1038/s41586-021-04072-3.

Yoav Benjamini and Yosef Hochberg. Controlling the false discovery rate: A practical and powerful approach to multiple testing. Journal of the Royal Statistical Society: Series B (Methodological), 57(1):289–300, 1995. doi: 10.1111/j.2517-6161.1995.tb02031.x.

Robert C. Edgar. Muscle5: high-accuracy alignment ensembles enable unbiased assessments of sequence homology and phylogeny. Nat Commun, 13:6968, 2022. doi: 10.1038/s41467-022-34630-w.

Zhuoqing Fang, Xinyuan Liu, and Gary Peltz. GSEApy: a comprehensive package for performing gene set enrichment analysis in python. Bioinformatics, 39(1):btac757, 2023. doi: 10.1093/bioinformatics/btac757.

Joseph Felsenstein. Phylogenies and the comparative method. Am Nat, 125(1):1–15, 1985. doi: 10.1086/284325.

Irene Gallego Romero, Ilya Ruvinsky, and Yoav Gilad. Comparative studies of gene expression and the evolution of gene regulation. Nat Rev Genet, 13(7):505–516, 2012. doi: 10.1038/nrg3229.

Jr. Garland Theodore and Anthony R. Ives. Using the past to predict the present: confidence intervals for regression equations in phylogenetic comparative methods. Am Nat, 155(3):346–364, 2000. doi: 10.1086/303327.

Peter W. Harrison and Stephen H. Montgomery. Genetics of cerebellar and neocortical expansion in anthropoid primates: A comparative approach. Brain Behav Evol, 89(4):274–285, 2017. doi: 10.1159/000477432.

Paul H. Harvey and Mark D. Pagel. The Comparative Method in Evolutionary Biology. Oxford University Press, Oxford, 1991.

Suzana Herculano-Houzel. Neuronal scaling rules for primate brains: the primate advantage. Prog Brain Res, 195:325–340, 2012. doi: 10.1016/B978-0-444-53860-4.00015-5.

Suzana Herculano-Houzel, Christine E. Collins, Peiyan Wong, and Jon H. Kaas. Cellular scaling rules for primate brains. Proc Natl Acad Sci U S A, 104(9):3562–3567, 2007. doi: 10.1073/pnas.0611396104.

Suzana Herculano-Houzel, Kamilla Avelino-de Souza, Kleber Neves, Jairo Porfírio, Débora Messeder, Larissa Mattos Feijó, José Maldonado and Paul R. Manger. The elephant brain in numbers. Front Neuroanat, 8:46, 2014. doi: 10.3389/fnana.2014.00046.

Suzana Herculano-Houzel, Kenneth Catania, Paul R. Manger, and Jon H. Kaas. Mammalian brains are made of these: a dataset of the numbers and densities of neuronal and nonneuronal cells in the brain of Glires, Primates, Scandentia, Eulipotyphlans, Afrotherians and Artiodactyls, and their relationship with body mass. Brain Behav Evol, 86(3-4): 145–163, 2015. doi: 10.1159/000437413.

Subha Kalyaanamoorthy, Bui Quang Minh, Thomas K. F. Wong, Arndt von Haeseler, and Lars S. Jermiin. ModelFinder: fast model selection for accurate phylogenetic estimates. Nat Methods, 14(6):587–589, 2017. doi: 10.1038/nmeth.4285.

Philipp Khaitovich, Wolfgang Enard, Michael Lachmann, and Svante Pääbo. Evolution of primate gene expression. Nat Rev Genet, 7(9):693–702, 2006. doi: 10.1038/nrg1940.

Amanda Kowalczyk, Wynn K. Meyer, Raghavendran Partha, Weiguang Mao, Nathan L. Clark, and Maria Chikina. RERconverge: an R package for associating evolutionary rates with convergent traits. Bioinformatics, 35(22):4815–4817, 2019. doi: 10.1093/bioinformatics/btz468.

Jovana Maliković, Juan L. Cantalapiedra, Lorenzo Vinciguerra, Katja Schönbächler, Ana Luiza F. Destro, Jennifer Rodger, Marielle Jörimann, Liora Las, Stephen G. Hörpel, David P. Wolfer, Lutz Slomianka, and Irmgard Amrein. Traces of phylogeny and ecology in hippocampal neuron numbers. PNAS Nexus, 4(9):pgaf261, 2025. doi: 10.1093/pnasnexus/pgaf261.

Sergey Melnikov, Adam Ben-Shem, Nicolas Garreau de Loubresse, Lasse Jenner, Gulnara Yusupova, and Marat Yusupov. One core, two shells: Bacterial and eukaryotic ribosomes. Nature Structural & Molecular Biology, 19(6): 560–567, 2012. doi: 10.1038/nsmb.2313.

Yoshihito Niimura and Masatoshi Nei. Extensive gains and losses of olfactory receptor genes in mammalian evolution. PLoS ONE, 2(8):e708, 2007. doi: 10.1371/journal.pone.0000708.

Raghavendran Partha, Amanda Kowalczyk, Nathan L. Clark, and Maria Chikina. Robust method for detecting convergent shifts in evolutionary rates. Molecular Biology and Evolution, 36(8):1817–1830, 2019. doi: 10.1093/molbev/msz107.

Antonio Pinto-Duarte, Katharine Bogue, Terrence J. Sejnowski, and Shyam Srinivasan. Conservation of glial density and cell-type ratios within a brain region across mammals. PNAS Nexus, 4(11):pgaf314, 2025. doi: 10.1093/pnasnexus/pgaf314.

Jeffrey Rogers and Richard A. Gibbs. Comparative primate genomics: emerging patterns of genome content and dynamics. Nat Rev Genet, 15(5):347–359, 2014. doi: 10.1038/nrg3707.

Diana K. Sarko, Kenneth C. Catania, Duncan B. Leitch, Jon H. Kaas, and Suzana Herculano-Houzel. Cellular scaling rules of insectivore brains. Front Neuroanat, 3:8, 2009. doi: 10.3389/neuro.05.008.2009.

Yong Shao, Long Zhou, Fang Li, Lan Zhao, Bao-Lin Zhang, Feng Shao, Jia-Wei Chen, Chun-Yan Chen, Xupeng Bi, Xiao-Lin Zhuang, Hong-Liang Zhu, Jiang Hu, Zongyi Sun, Xin Li, Depeng Wang, Iker Rivas-González, Sheng Wang, Yun-Mei Wang, Wu Chen, Gang Li, Hui-Meng Lu, Yang Liu, Lukas F. K. Kuderna, Kyle Kai-How Farh, Peng-Fei Fan, Li Yu, Ming Li, Zhi-Jin Liu, George P. Tiley, Anne D. Yoder, Christian Roos, Takashi Hayakawa, Tomas Marques-Bonet, Jeffrey Rogers, Peter D. Stenson, David N. Cooper, Mikkel Heide Schierup, Yong-Gang Yao, Ya-Ping Zhang, Wen Wang, Xiao-Guang Qi, Guojie Zhang, and Dong-Dong Wu. Phylogenomic analyses provide insights into primate evolution. Science, 380 (6648):913–924, 2023. doi: 10.1126/science.abn6919.

Chet C. Sherwood, Sarah B. Miller, Molly Karl, Cheryl D. Stimpson, Kimberley A. Phillips, Bob Jacobs, Patrick R. Hof, Mary Ann Raghanti, and Jeroen B. Smaers. Invariant synapse density and neuronal connectivity scaling in primate neocortical evolution. Cerebral Cortex, 30(10):5604–5615, 2020. doi: 10.1093/cercor/bhaa149.

Mehmet Somel, Xiling Liu, and Philipp Khaitovich. Human brain evolution: transcripts, metabolites and their regulators. Nat Rev Neurosci, 14(2):112–127, 2013. doi: 10.1038/nrn3372.

Fredrik Tegenfeldt, Dmitry Kuznetsov, Mosè Manni Matthew Berkeley, Evgeny M. Zdobnov, and Evgenia V. Kriventseva. OrthoDB and BUSCO update: annotation of orthologs with wider sampling of genomes. Nucleic Acids Res, 53(D1):D516–D522, 2025. doi: 10.1093/nar/gkae987.

Fredj Tekaia. Inferring orthologs: Open questions and perspectives. Genomics Insights, 9:17–28, 2016. doi: 10.4137/GEI.S37925.

Nathan S. Upham, Jacob A. Esselstyn, and Walter Jetz. Inferring the mammal tree: species-level sets of phylogenies for questions in ecology, evolution, and conservation. PLoS Biol, 17(12):e3000494, 2019. doi: 10.1371/journal.pbio.3000494.

Martijn P. van den Heuvel, Edward T. Bullmore, and Olaf Sporns. Comparative connectomics. Trends in Cognitive Sciences, 20(5):345–361, 2016. doi: 10.1016/j.tics.2016.03.001.

Pauli Virtanen, Ralf Gommers, Travis E. Oliphant, Matt Haberland, Tyler Reddy, David Cournapeau, Evgeni Burovski, Pearu Peterson, Warren Weckesser, Jonathan Bright, Stéfan J. van der Walt, Matthew Brett, Joshua Wilson, K. Jarrod Millman, Nikolay Mayorov, Andrew R. J. Nelson, Eric Jones, Robert Kern, Eric Larson, C. J. Carey, İlhan Polat, Yu Feng, Eric W. Moore, Jake VanderPlas, Denis Laxalde, Josef Perktold, Robert Cimrman, Ian Henriksen, E. A. Quintero, Charles R. Harris, Anne M. Archibald, Antônio H. Ribeiro, Fabian Pedregosa, Paul van Mulbregt, and SciPy 1.0 Contributors. SciPy 1.0: Fundamental algorithms for scientific computing in Python. Nature Methods, 17(3):261–272, 2020. doi: 10.1038/s41592-019-0686-2.

Cindy L. Will and Reinhard Lührmann. Spliceosome structure and function. Cold Spring Harbor Perspectives in Biology, 3 (7):a003707, 2011. doi: 10.1101/cshperspect.a003707.

Thomas K. F. Wong, Nhan Ly-Trong, Huaiyan Ren, Piyumal Demotte, Hector Baños, Andrew J. Roger, Edward Susko, Chris Bielow, Nicola De Maio, Nick Goldman, Matthew W. Hahn, Mario Dos Reis, Le Sy Vinh, Gavin Huttley, Robert Lanfear, and Bui Quang Minh. IQ-TREE 3: phylogenomic inference software using complex evolutionary models. Mol Biol Evol, 43(5):msag117, 2026. doi: 10.1093/molbev/msag117.

Linhe Xu and Suzana Herculano-Houzel. Neuron-specific homologous coding genes and non-coding regulatory regions are the most conserved amongst amniotes despite neuron-specific cell size diversity. Scientific Reports, 15(1):43971, 2025. doi: 10.1038/s41598-025-27680-9.

Xiao-Lin Zhuang, Jin-Jin Zhang, Yong Shao, Yaxin Ye, Chun-Yan Chen, Long Zhou, Zheng-Bo Wang, Xing-Yan Zhang, and Dong-Dong Wu. Integrative omics reveals rapidly evolving regulatory sequences driving primate brain evolution. Molecular Biology and Evolution, 40(8):msad173, 2023. doi: 10.1093/molbev/msad173.

